# Fast draft assemblies using locally consistent parsing

**DOI:** 10.64898/2026.11.23.768163

**Authors:** Fatih Karaoğlanoğlu, Klea Zambaku, Akmuhammet Ashyralyyev, Ezgi Ebren, Lucia Nappi, Faraz Hach, Can Alkan

## Abstract

Recent improvements in long-read sequencing accuracy, particularly the introduction of PacBio HiFi reads, have created new opportunities for more efficient sequence representations in genome assembly. In particular, variations of de Bruijn graphs (dBG), such as sparse dBG and minimizer-space dBG, provide efficient representations of sequences and have enabled genome assemblers to leverage the high accuracy of long-read data while maintaining low memory usage and fast analysis times. Locally consistent parsing (LCP) is another technique for compact sequence representation that partitions a string into nearly uniform, highly distinctive substrings in a consistent manner. In this work, we combine ideas from minimizer-space assembly and LCP to achieve a significant speedup in HiFi long-read assembly. Specifically, we define a locally consistent de Bruijn Graph (lcDBG), in which string partitions serve as the graph’s building blocks. We introduce Gallifrey, a rapid PacBio HiFi assembler that outperforms the fastest existing tool, rust-mdbg, by *>* 6× in runtime while requiring 2×less memory, assembling a human genome in under one minute on a standard laptop, while generating assemblies with comparable contiguity. Code for Gallifrey is freely available at https://github.com/BilkentCompGen/gallifrey.

## 1 Introduction

Two decades after their introduction, high-throughput sequencing (HTS) [1] technologies have become the primary driving force across all research areas in the life sciences. Recent developments in sequencing technologies have improved both the accuracy and the read lengths. Furthermore, with consensus sequencing techniques such as the PacBio HiFi platform, it has become possible to obtain highly accurate long reads [2]. This led to the development of new assemblers using long reads, such as Flye [3], and those specifically designed for HiFi data, such as HiCanu [4] and hifiasm [5], which generated more complete and accurate assemblies. Later, the Verkko assembler [6] was developed as a hybrid workflow that combined HiCanu [4], MBG [7], and GraphAligner [8] to generate Telomere-to-Telomere assemblies [9] using HiFi reads together with longer, more error-prone Oxford Nanopore reads. More recently, Verkko2 was developed to enhance Verkko’s computational efficiency and also introduced improved haplotype phasing, scaffolding, and gap closing [10].

There are three popular graph structures, including their derivatives and hybrids, used for genome assembly [11]. *Overlap-Layout-Consensus (OLC) graphs* represent each sequence read as a node, and connect nodes with edges that indicate a significant overlap between them. These overlaps are found using computationally expensive mapping and alignment procedures. *De Bruijn graphs* represent all *k*-mers (substrings of length *k*) from the reads as nodes and connect two nodes if they overlap by *k*− 1 bases within reads. Finally, *string graphs*, which are modifications of OLC graphs, represent reads as nodes and connect them with edges only if they have an irreducible overlap (one not already contained within a longer overlap path).

De Bruijn graphs further partition reads into smaller segments of length *k* (i.e., *k*-mers), which can yield highly cyclic and fragmented graphs. Despite this shortcoming, they also offer several advantages: 1) they do not require all-by-all pairwise alignments, reducing the amount of computation required; 2) reads can be processed independently of one another, allowing for better parallelization; and 3) graph size (i.e., number of *k*-mers) scales linearly with the genome size; however, erroneous *k*-mers generated because of sequencing errors increase memory usage. Therefore, it is still common to use de Bruijn graphs and their derivatives, such as A-Bruijn graphs [12] and multiplex de Bruijn graphs [13], for genome assembly, particularly with highly accurate long reads (i.e., PacBio HiFi). The traditional de Bruijn graph (dBG) construction uses all *k*-mers in the input data, which can consume significant memory when sequencing errors are present. Furthermore, traditional methods are not able to increase the size of *k*-mers to fully exploit the benefits of long reads. To address these problems, alternative dBG constructions are developed, which can be exemplified by MBG [7] and rust-mdbg [14]. These sketching-based approaches [15] use a sparse subset of *k*-mers (i.e., minimizers) rather than the full *k*-mer spectrum. This fundamental shift to a sparse representation results in a graph with dramatically fewer nodes and edges, substantially reducing memory requirements and accelerating construction. Building upon sparse de Bruijn graph principles, rust-mdbg introduces the minimizer-space DBG, wherein the fundamental units of the graph are minimizers rather than nucleotide bases. This sparsity also provides implicit graph compaction, as non-branching paths between minimizers are condensed, simplifying the graph topology from the outset. rust-mdbg was later extended to metaMDBG [16] to assemble metagenomes, which typically exhibit variable genome coverage.

While rust-mdbg enables rapid assembly graph construction, flat sketching schemes face a fundamental trade-off between sparsity, positional variance, and context sensitivity. Universal minimizers select *k*-mers independently via a hash threshold, yielding geometrically distributed inter-seed distances with high variance (CV *≈* 1 under independent hashing [14,17]) that either leave long unsketched gaps or require high sampling density [14,18]. Windowed minimizers bound the maximum gap but suffer from context sensitivity, where a single mutation near a window boundary can shift neighboring minimizer selections. Conversely, while iterative Locally Consistent Parsing (LCP) [19,20,21] guarantees bounded-overlap string partitioning—achieving a near-uniform intercore distance coefficient of variation of CV *≈* 0.28 at higher levels on human assemblies [21], applying nucleotide-level LCP directly to 30× human HiFi reads requires constructing and reducing over 4 × 10^10^ level-1 cores (Supplementary Section S2), shifting the bottleneck to base-level parsing [21]. Furthermore, unlike canonical *k*-mers that symmetrically collapse forward and reverse-complement strands, Deterministic Coin Tossing (DCT) [22] and lexicographical LCP rules are inherently directional (chiral): a DNA sequence and its reverse complement produce distinct core partitions, preventing naive double-stranded graph construction.

Here, we present Gallifrey, a de Bruijn graph-based long-read assembler that decouples hierarchical sparsity from base-level parsing overhead while resolving strand asymmetry in locally consistent graphs. Building upon our iterative LCP framework (Lcptools) [21], Gallifrey introduces four algorithmic contributions: (i) **Locally Consistent Seed Clouds (LCSC):** Gallifrey anchors level-1 parsing on context-free open syncmers [23] and iteratively applies DCT and LCP rules to seed vectors (Section 2.2). This bypasses the *O*(*n*) base-level DCT bottleneck, computes all higher hierarchy levels in *O*(*m*_1_) total time (where *m*_1_ = Θ (*n/*(*k* − *s* + 1))), and achieves an expected ∼ 270× sparsification relative to homopolymer-compressed bases (Supplementary Section S2, Proposition S1). A single-character edit perturbs at most a constant number of cores at any depth (Lemma 1). (ii) **Constant-Time Sketch-Space Encoding and Delta Storage:** Gallifrey represents graph vertices as *W* -core windows (*lcmers*) packed into collision-free 64-bit hybrid keys (combining the exact 32-bit identifier of the terminal core with a 32-bit Karp–Rabin polynomial rolling hash [24] of the *W* − 1 prefix cores; Section 2.4, Algorithm S1), paired with nonredundant core-suffix delta sequence storage (Section 2.5) for single-pass nucleotide reconstruction. (iii) **Chiral Strand Reconciliation via Retro Admission and Core-Space DAG Consensus:** To address the directional asymmetry of LCP in double-stranded DNA, Gallifrey combines a lock-free blocked Bloom filter gate [25] with reverse-complement priming and retroactive edge admission during streaming construction. This reduces expected single-strand coverage dropouts by over an order of magnitude (Equations 2–3). A subsequent bipartite interval-overlap layout and core-space DAG consensus (Section 2.7) bridge staggered single-strand breakpoints. (iv) **Lock-Free Parallel Path Contraction:** Gallifrey partions maximal simple paths into mutually exclusive, node-disjoint chains uniquely anchored at terminal nodes (Section S3, Algorithm S6), enabling lock-free and race-free parallel unitig contraction without synchronization overhead across sharded concurrent hash tables. Across *D. melanogaster* and human CHM13 PacBio HiFi datasets, Gallifrey ran 2.5× − 6.1× faster and used 2× −3 × less memory than rust-mdbg, assembling a human genome in under one minute on a laptop while reducing misassemblies by an order of magnitude.

## 2 Methods

GALLIFREY consists of three stages: graph building, graph simplification, and reverse-complement folding. In the graph-building stage, multiple worker threads compute lcmers and their adjacencies and then insert them into a sharded hash table representing the de Bruijn graph. In the graph-simplification stage, GALLIFREY iteratively constructs unitigs by simplifying the graph and removing tips. In the reverse-complement folding stage, a bipartite overlap graph is built to connect unitigs with their reverse-complement overlaps. By finding and combining the connected components in this overlap graph, GALLIFREY merges reverse-complementary unitigs and improves the assembly contiguity.

### 2.1 Locally consistent parsing

Locally Consistent Parsing (LCP) is a string-partitioning technique that constructs an overlapping set of near-uniform-length substrings, called *cores*, which are distributed uniformly across the string. The main idea of LCP is to process the string hierarchically, first identifying cores at the lowest level and then iteratively combining them to form more compact cores. Hence, as the iterative processing continues, all properties are propagated to the upper-level cores, yielding a significantly smaller set of compact substrings that represent the original string. LCP constructs cores if the substrings are aligned with one of the four patterns based on lexicographical ordering, which are *local minima* (LMIN), *local maxima* (LMAX), *repetitive characters* (RINT), or *stranded sequences* (SSEQ). The last type of core closes any gap among the first three cases. After constructing the first-level cores, the DCT is applied to the cores’ identifiers, referred to as bitstreams [21], to reduce the alphabet size and maintain the desired partitioning and uniformity properties [22,19] once the new cores are constructed.

### 2.2 Locally Consistent Seed Cloud

We introduce a new spaced-seeding technique, LCSC, that applies a locally consistent parsing to vectors of seed sequences rather than directly to raw nucleotide strings. Given an input DNA sequence, we first apply homopolymer compression (HPC) to reduce the errors caused by homopolymer run-length variation (Section 2.3). From this compressed sequence, we select a seed vector that serves as input to the LCSC construction.

Each selected seed forms a level-1 LCSC core, the base layer of the hierarchy. Higher-level cores are then built iteratively, as in LCP (Figure 1.a): at each level, we apply DCT to the current sequence of cores, followed by the standard LCP rules, to identify the cores of the next level. This partitions each seed vector into consistent segments with stable identifiers, which we call *seed-cores*. The resulting hierarchy of seed-cores is the building block for downstream applications; in particular, Gallifrey constructs a de Bruijn graph whose alphabet is the seed-cores.

**Fig 1.**
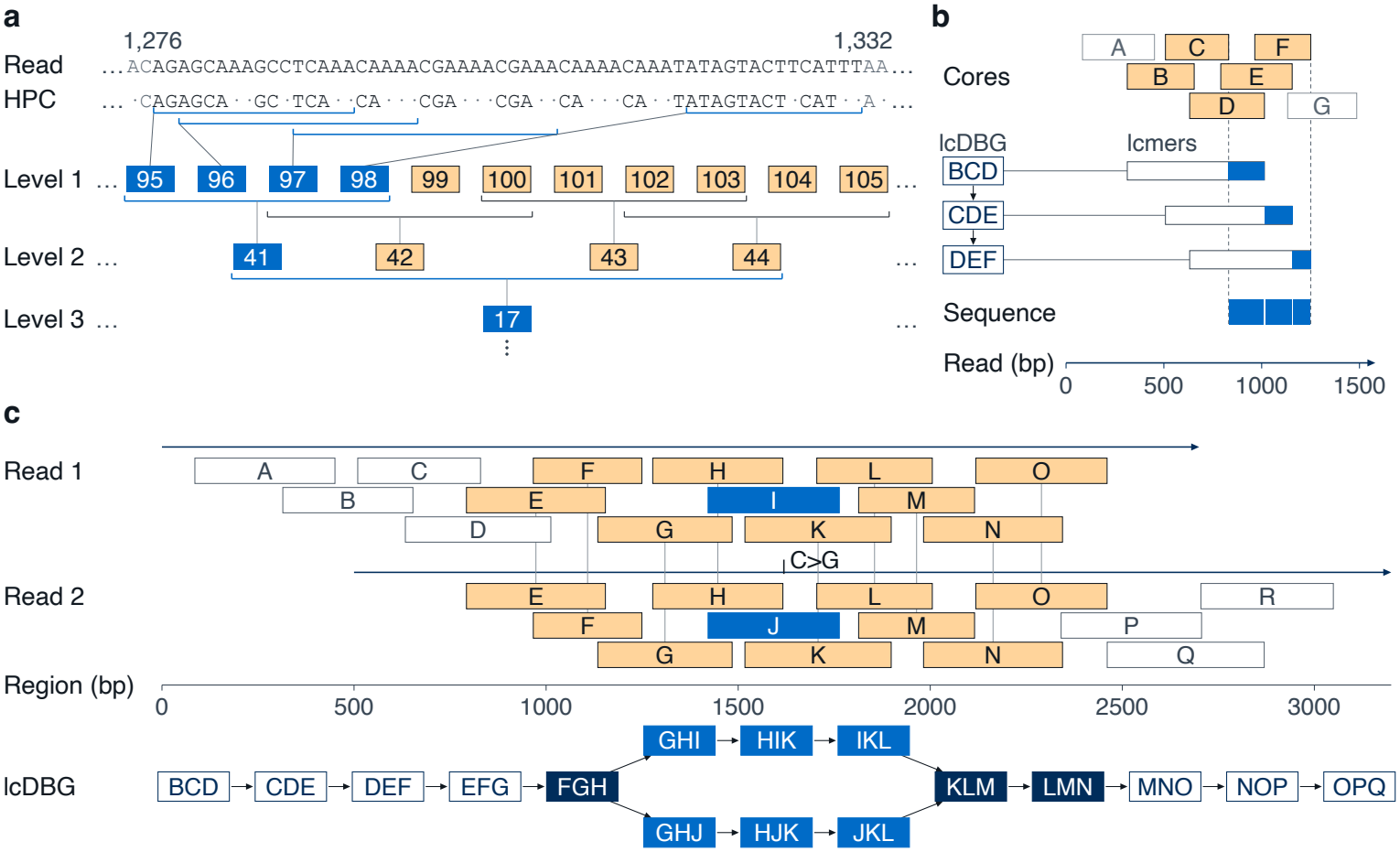
LCSC and the lcDBG. (a) Seeds selected from a homopolymer-compressed read form the level-1 LCSC cores, and higher-level cores are constructed iteratively from the cores of the previous level. (b) *W* consecutive cores (*W* = 3 here) form an lcmer, a node of the lcDBG. Each node stores only the sequence between the end positions of its last two cores, so that consecutive fragments along a path do not overlap and their concatenation reconstructs the genome sequence without redundancy. (c) Because the parsing is locally consistent, two overlapping reads yield the same cores in their shared region, and a substitution changes only the core that contains it. The lcmer’s containing this core form a bubble in the lcDBG.

Furthermore, LCSC can be viewed as a generalization of LCP: when *k* = 1, such that fixed-length seeds of length 1 are used, and all possible seeds (*A, C, G*, and *T*) are valid, LCSC is equivalent to LCP. LCSC also can be viewed as a subsampling scheme over the seeds of strobemers [18] (with a variable number of strobes), while providing LCP guarantees of sequence coverage and predictable overlaps between adjacent units.

LCSC is designed to work with any consistent, context-independent seed selection method that does not rely on surrounding sequences. Gallifrey uses open syncmers [23] with parameters (*k, s, t*), in which a *k*-mer is selected as a seed if the smallest of its *s*-mers lies at offset *t*. Because seed selection is context-independent and each level of LCP inspects only a constant-size neighborhood, a sequence edit perturbs only a constant number of seed-cores at every level, and only a bounded bubble in the lcDBG (Lemma 1).

#### Lemma 1

**(Locality of perturbations)**.

*Let* 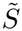 *and* 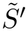 *be homopolymer-compressed strings that differ by a single-character edit, and let Δ* ^*ℓ*^ *denote the length of the differing block of seed-cores at level `. Assume open syncmers* (*k, s, t*) *and let d* = min(*t, k − s − t*) + 1.

1. Level 1: *Δ*_*1*_ ≤ *⌈dk/de⌉, and each differing block has expected size at most k/*(*k−s*+1). *For k* = 12, *s* = 4, *t* = 5, *this gives Δ*_*1*_ ≤ 3.
2. Higher levels: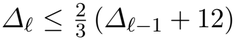, *where the constant* 12, *independent of the seed parameters, accounts for the neighbors that DCT and the LCP rules inspect, and* 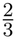*follows from the Adjacency Property (i*.*e*., *worst-case bound for* 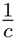 *[21]). Because Δ*_*1*_ ≤3, *this gives Δ*_*1*_ ≤ 24 *for every ℓ, independent of read length and hierarchy depth. With the empirical reduction factor c ≈* 2.34 *[21], the bound drops to ∼* 9.
3. lcDBG: *the edit changes at most Δ* _*L*_ + *W −* 1 *consecutive lcmers, so it forms a bubble with arms of at most Δ* _*L*_ + *W −* 1 *nodes. For Δ* _*L*_ = 1, *the arms have exactly W nodes (Fig. 1c)*.

*Proofs are given in Supplementary Section S2*.

### 2.3 Homopolymer reduction

Homopolymer compression (HPC) is now a standard technique used by many mapping and assembly methods [26,14,7], in which homopolymer sequences are collapsed into single characters and their run lengths are stored for later reconstruction.

Gallifrey similarly applies HPC to compute seed positions, core boundaries, and core identifiers without storing per-base run-length arrays during parsing. However, Gallifrey maps the resulting core boundaries back to the original read coordinates and stores the uncompressed nucleotide substrings inside graph nodes (Section 2.5), thereby assembling full-length, non-compressed genomic sequences.

### 2.4 Graph construction

The primitives of the lcDBG under construction are runs of *W* consecutive LCSC cores, hereafter referred to as *lcmers* (Figure 1.b); the construction procedure is outlined in Algorithm S2. These lcmers are encoded as 64-bit integers, where the lower 32 bits store the last LCSC core in the lcmer window, and the upper 32 bits store a Karp–Rabin-style polynomial rolling hash [24] of the *W* − 1 previous cores (Algorithm S1). In each step, the oldest core multiplied by *B*^*W*^ − ^2^ is subtracted from the upper bits, which are then multiplied by an odd constant *B* and incremented by the lower bits, and the lower bits are replaced by the new core. This scheme allows us to efficiently compute and store windows of LCSC cores while preserving their order information. In our experiments, we observed no collisions between lcmers during the assembly of the human genome. To reduce the number of artifactual lcmer’s created by sequencing errors, an lcmer is admitted to the graph only from its second sighting onward: a blocked Bloom filter [27,25] records every sighting, and an lcmer whose key the filter does not yet hold is skipped. An admitted lcmer creates/updates its node only if a neighboring lcmer of the same read is also admitted. If the preceding lcmer is admitted, an edge from it to this lcmer is added; if the following one is admitted, this node’s count is incremented. Using a thread-safe hash table and Bloom filter, the graph can be constructed in parallel.

#### lcmer coverage

The gate admits an lcmer only after two sightings. A read can sight an lcmer only if it contains the whole window and no error in the window changes its seed-cores. Assuming reads are longer than windows and treating read starts as a Poisson process, the number of such reads at window *w* is Poisson with mean

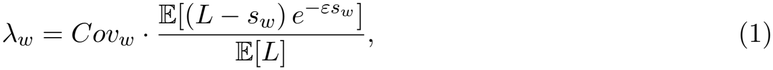

where *Cov*_*w*_ is the local read coverage, *L* a read’s length, *s*_*w*_ the window’s span and *ε* a read’s rate of errors visible after homopolymer compression.

#### Gate dropouts, priming, and retro admission

A window not admitted to the graph leaves a strand break. Under the two-sighting gate alone, an lcmer requires two reads on its own strand, leaving a coverage gap with probability:

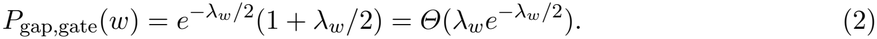

To mitigate strand dropouts, Gallifrey applies *reverse-complement priming* by recording observed complementary windows under a strand-tagged key (Algorithm S2), reducing gap probability To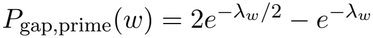.

Furthermore, Gallifrey introduces *retro admission*: subsequent reads on the complementary strand can instantiate a dropped forward node and its incident edges (Algorithm S2; Supplementary Section S3). Together, priming and retro admission yield:

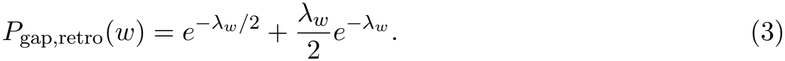

Equation (3) matches the physical coverage limit of a streaming, strand-chiral de Bruijn graph: a locus fails admission only if it lacks reads on the forward strand, or has exactly one forward read and zero complementary reads. In typical 33*×* HiFi human datasets (*λ*_*w*_ *≈* 20) [28,4], this reduces expected window dropouts by over an order of magnitude, from *∼* 8,000 down to *∼* 700 (Supplementary Section S1.1).

These estimations assume uniform read coverage (Equation 1). This is true for most of the genome with HiFi reads, except in long A/G- or C/T-rich, low-complexity sequence regions where read coverage drops considerably. This introduces another set of coverage gaps that are inherent to the sequencing protocol and can only be avoided by bridging the assembly with reads from other protocols (e.g., nanopore).

### 2.5 Storing sequences

By construction, consecutive LCP cores overlap, and this also holds for LCSC cores. A core may overlap each of its preceding cores by a different amount, but the overlap between two given cores is always the same. When a new lcmer *ρ* [*i − W* + 1 ... *i*] is added to the graph, Gallifrey stores in its node the difference between its last two cores, that is, the sequence from the end of *ρ* [*i−* 1] to the end of *ρ* [*i*]. The assembled sequences are then generated by traversing the graph and appending the stored sequences. This stores some sequence more than once: a core that is the last core of several different lcmers is stored in each of them. In return, it preserves the differences between sequences that share the same *k*-mer seed representation but differ in homopolymer lengths or in bases not covered by syncmers, as long as they lie in different lcmers.

### 2.6 Graph simplification and cleaning

After construction, Gallifrey simplifies the lcDBG in rounds until it converges (Supplementary Section S3). Each round merges non-branching paths into unitigs, removes tips, and pops bubbles. Because most nodes are simple after construction, the first merge is the most expensive step, and we parallelize it without locks. Every maximal simple path has a unique last node, which can be recognized from that node and its immediate neighbors alone. All paths can therefore be found in parallel, and since they are node-disjoint, they can be merged concurrently with no shard locks and no races.

The cleaning thresholds are derived from the node-depth distribution. The one-copy depth *d*_1_ is the lowest substantial peak of the depth distribution of long nodes, and the tip depth is *d*_1_*/*2. A bubble is popped only if its removed arm is short relative to the heaviest path and none of its nodes reaches the locus depth limit *d*_max_, because deeper bubbles usually arise from repeat copies rather than sequencing errors. The same limit *d*_max_ is reused for dead-end joining (Section 2.8). We provide further details of graph simplification in Section S3.

### 2.7 Bi-partite overlap-layout

LCP and LCSC are strand-specific. Due to their directional construction, a string and its reverse complement yield different partitions, even when the seeds are selected in the same way on both strands. Complementary sequences therefore cannot be merged during graph construction, and a locus is usually represented by unitigs of both strands, which may break at different places (Figure 2.a). This requires Gallifrey to merge complementary unitigs after graph cleaning.

**Fig 2.**
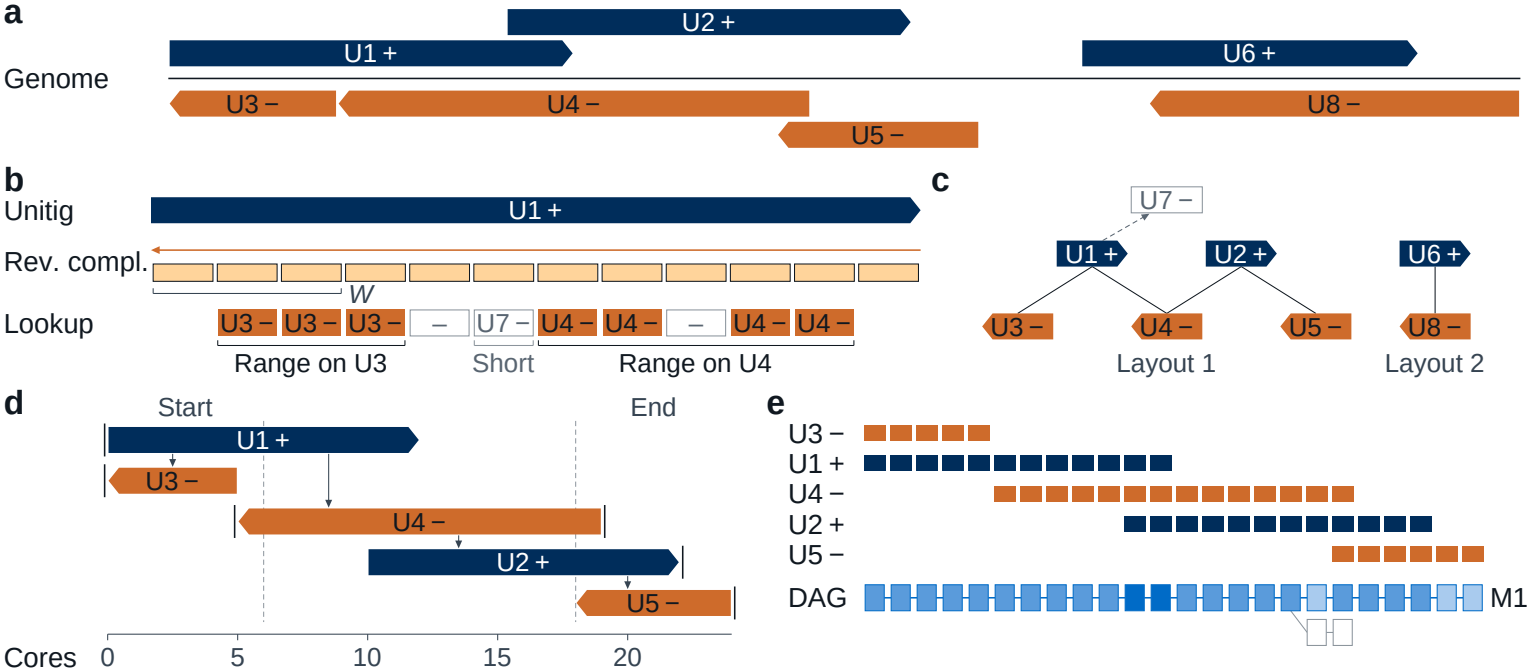
Strand reconciliation and layout. (a) Unitigs of the two strands along the genome. If unitigs end at different breakpoints, we can merge them into a single sequence in this step. (b) lcmer’s of each unitig’s reverse complement sequences are looked up in the cleaned lcDBG, and overlapping intervals between each unitig pair are determined. (c) The overlap graph is constructed by connecting pairs of unitigs with reciprocal overlaps, indicating that they represent equivalent sequences in opposite orientations. Each connected component in this overlap graph is called a layout. (d) Doing breadth-first iteration, we assign +signs and core positions relative to other unitigs in the layout. We preserve the edges of the unitigs on the boundaries and connected to the newly created layout node. (e) The unitigs are merged in the order of their layouts into a DAG based on LCSC cores, where nodes are matched and collapsed together if their cores share the same label and HPC sequence. The heaviest path of each weakly connected component of the DAG becomes a segment.

For each unitig, we compute the LCSC cores of its reverse complement and look up its lcmers in the cleaned graph, where every lcmer leads to the unitig that contains it (Figure 2.b). Consecutive hits on the same unitig form an overlap range: an lcmer that is not found does not end the range, while a hit on another unitig starts a new one. An overlap between two unitigs is kept only if it is reciprocal, that is, the reverse complement of each unitig hits the other, and if it spans at least max(2, ⌊*W/*2 ⌋) cores on both unitigs. We then construct an undirected bipartite overlap graph, in which each node represents a unitig and each edge indicates that the corresponding pair of unitigs overlap as reverse complements (Figure 2.c).

In this graph, a connected component can be one of three types: a singleton, meaning no reverse complement match was found; a one-to-one match, indicating that the forward and reverse orientations break at the same places, typically due to sequence duplication in the genome; or a larger component, in which breakpoints are not shared between orientations. In the latter case, we combine the unitigs using the linking information derived from the complementary sequences, thereby improving assembly contiguity.

For each connected component, we first compute a layout by assigning relative positions and orientations from the pairwise overlaps (Figure 2.d). This is done with BFS: the first unitig is assigned position 0 and the forward orientation; each edge flips the orientation, and relative positions, counted in cores, follow from the overlap ranges of the matched unitigs. Unitigs that start or end near an end of the layout are marked as its boundaries.

We then merge the unitigs of each layout into a DAG of LCSC cores (Figure 2.e, Algorithm S5). Each unitig is parsed again in its assigned orientation and added in order of position. A core is merged into an existing node only if both have the same label and the same homopolymer-compressed sequence, and only if the match belongs to a chain of matches that agrees with the pseudo-layout. Isolated matches, typically cores shared with another copy of a repeat, are discarded. The weight of a node is the number of unitigs that share it, and the heaviest path of each weakly connected component of the DAG becomes a segment. A layout with at most one unitig per orientation skips the DAG, and its longer unitig becomes the segment.

Finally, we replace the unitigs of each layout with its segment. Edges incident to the boundary unitigs of a layout are redirected to the end of the segment that matches the unitig’s orientation, and edges incident to the other unitigs of the layout are removed. Unitigs outside layouts whose only connections lead into the interiors of layouts are removed as well, as they are alternatives to sequence already inside a layout. Short segments outside layouts that have links on only one side are then removed repeatedly until none remain; layout segments are never removed.

### 2.8 Post-layout dead-end joining

The layout merges two strands only where their unitigs overlap by enough lcmers to be detected (Section 2.7). When both strands break close to each other, this overlap is too short, and the locus remains split into two segments. Each ends in a dead end, a segment side without links, and the two dead ends overlap as reverse complements.

To find these overlaps, we compute the LCSC cores of the last bases of every dead end one level below the assembly level, so that even short overlaps contain several cores. Two dead ends that share cores in a consistent order are aligned with the banded extension alignment of minimap2 [26], and the overlap is accepted if the alignment reaches both ends at high identity. A pair is joined only if neither end overlaps any other end, both ends lie below the locus depth limit *d*_max_, and neither end lies in a tandem repeat, where the join point would be ambiguous. Each chain of joined segments becomes one segment, and ends are never joined across gaps.

## 3 Results

We evaluated Gallifrey on PacBio HiFi reads from two genomes differing in scale and repeat content: fruit fly (*D. melanogaster*) and the haploid human cell line CHM13. We compared it with rust-mdbg, the fastest existing HiFi assembler and the most similar in design, since both build a sparse de Bruijn graph in sketch space. We report runtime and peak memory usage across thread counts, along with assembly contiguity and correctness measured using QUAST [29] against the corresponding reference genomes. For reference, we also include Hifiasm [5] and HiCanu [4]. Both are full-featured assemblers that perform read correction and consensus, which Gallifrey and rust-mdbg do not. They are therefore considerably slower but produce more contiguous assemblies and are standard practice. They serve as reference points for assembly contiguity rather than as direct competitors. We ran all tools on a server (parameters: Table S1) and additionally evaluated Gallifrey and rust-mdbg on a laptop to assess whether a human genome can be assembled on commodity hardware.

### 3.1 Performance analysis on a server node

#### Runtime and peak memory

We ran all benchmarks on a dual-socket server with two Intel Xeon Gold 6330N processors (2.20 GHz, 56 physical cores, 112 logical threads, 768 GB RAM). We evaluated Gallifrey on two genomes. The first comprised PacBio HiFi reads from SRR10238607 at 100*×* coverage for *D. melanogaster*. The second comprised CHM13 PacBio HiFi reads of *∼*10 Kbp (SRR9087597–SRR9087600) and *∼*20 Kbp (SRR11292120–SRR11292123), combined into a single dataset at 58*×* coverage. We ran both Gallifrey and rust-mdbg on both datasets. Table 1 reports their runtime and peak memory usage with 1 to 64 threads.

**Table 1.**
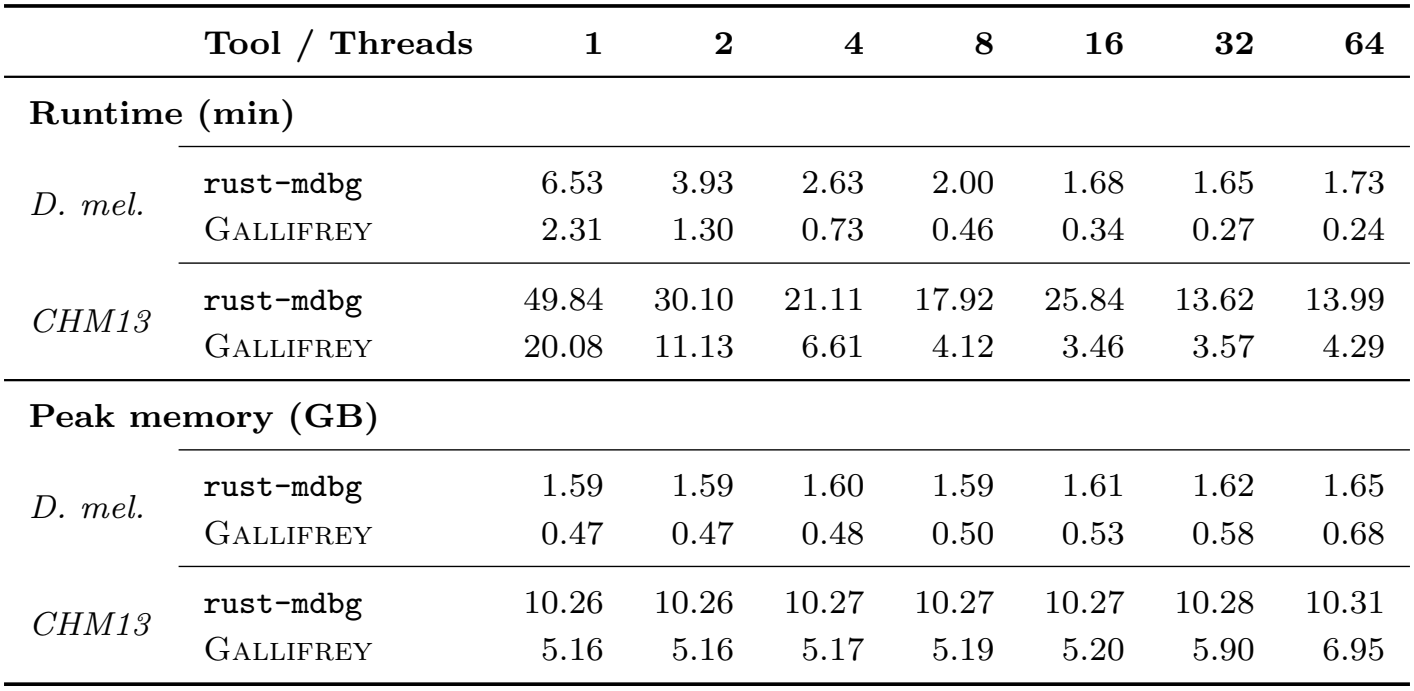
Runtime (minutes) and peak memory usage (GB) by thread count.

|  | **Tool / Threads** | **1** | **2** | **4** | **8** | **16** | **32** | **64** |
|---|---|---|---|---|---|---|---|---|
| **Runtime (min)** |  |  |  |  |  |  |  |  |
| *D. mel.* | rust-mdbg | 6.53 | 3.93 | 2.63 | 2.00 | 1.68 | 1.65 | 1.73 |
|  | Gallifrey | 2.31 | 1.30 | 0.73 | 0.46 | 0.34 | 0.27 | 0.24 |
| *CHM13* | rust-mdbg | 49.84 | 30.10 | 21.11 | 17.92 | 25.84 | 13.62 | 13.99 |
|  | Gallifrey | 20.08 | 11.13 | 6.61 | 4.12 | 3.46 | 3.57 | 4.29 |
| **Peak memory (GB)** |  |  |  |  |  |  |  |  |
| *D. mel.* | rust-mdbg | 1.59 | 1.59 | 1.60 | 1.59 | 1.61 | 1.62 | 1.65 |
|  | Gallifrey | 0.47 | 0.47 | 0.48 | 0.50 | 0.53 | 0.58 | 0.68 |
| *CHM13* | rust-mdbg | 10.26 | 10.26 | 10.27 | 10.27 | 10.27 | 10.28 | 10.31 |
|  | Gallifrey | 5.16 | 5.16 | 5.17 | 5.19 | 5.20 | 5.90 | 6.95 |

#### Runtime

On *D. melanogaster*, Gallifrey outperforms rust-mdbg at every thread count, achieving a 9.6*×* internal speedup from 1 to 64 threads versus *∼*4*×* for rust-mdbg, indicating more effective use of additional threads. At 1 thread, Gallifrey is 2.8*×* faster than rust-mdbg, increasing to 6.1*×* at 32 threads (rust-mdbg’s best) as Gallifrey scales more efficiently.

On CHM13, Gallifrey also outperforms rust-mdbg, which exhibits irregular scaling behaviour: runtime increases from 8 to 16 threads before decreasing at 32 threads, suggesting sensitivity to thread count on larger datasets. Gallifrey achieves a 5.8*×* internal speedup from 1 to 16 threads, versus *∼*2.4*×* for rust-mdbg from 1 to 32 threads. At rust-mdbg ‘s worst performance (1 thread), Gallifrey is 2.5*×* faster; at rust-mdbg ‘s best (32 threads), it is 3.8*×* faster. Across both datasets, Gallifrey achieves a 4.9*×* to 7.5*×* speedup over rust-mdbg at 16 threads.

#### Peak memory

On *D. melanogaster*, Gallifrey uses roughly 3*×* less memory than rust-mdbg at 16 threads (0.53 GB vs. 1.61 GB), with a modest increase at higher thread counts consistent with parallel workload partitioning. In contrast, rust-mdbg’s memory usage remains flat across all thread counts. On CHM13, the pattern is similar. Gallifrey uses approximately half the memory of rust-mdbg at 16 threads (5.20 GB vs. 10.27 GB). Although Gallifrey’s memory usage grows slightly at 64 threads, it remains well below rust-mdbg’s throughout.

#### QUAST contiguity analysis

For the evaluation, we used the dm6 assembly as the reference genome for *D. melanogaster* and CHM13v2.0 for the human genome experiments. Table 2 reports QUAST assembly quality metrics for *D. melanogaster* and *H. sapiens* (CHM13). We focus on metrics of contiguity and accuracy. N50 is the contig length such that contigs of this length or longer account for at least half the total assembly length; NG50 uses half the reference genome length instead, making it more meaningful for cross-assembly comparisons. NA50 and NGA50 are the alignment-aware counterparts of N50 and NG50, computed from aligned blocks obtained by breaking contigs at misassembly points, reflecting contiguity with respect to the reference.

**Table 2.** QUAST assembly quality metrics for *D. melanogaster* and *H. sapiens* (CHM13).

| **Genome** | **Metric** | **Hifiasm** | **HiCanu** | rust-mdbg | Gallifrey | Gallifrey-**MP**<sup>\*</sup> |
|---|---|---|---|---|---|---|
| *D. mel.* | N50 | 7,380,037 | 8,147,299 | 3,609,182 | 1,564,874 | — |
|  | NG50 | 22,069,487 | 18,283,070 | 8,327,409 | 3,331,917 | — |
|  | NA50 | 675,192 | 583,606 | 1,056,503 | 521,659 | — |
|  | NGA50 | 11,357,968 | 10,985,678 | 2,610,564 | 1,372,172 | — |
|  | # Misassemblies | 4,898 | 5,291 | 4,538 | 1,808 | — |
|  | Genome fraction (%) | 95.351 | 94.332 | 91.869 | 93.323 | — |
| CHM13 | N50 | 89,582,900 | 67,916,532 | 25,418,047 | 22,128,410 | 28,254,983 |
|  | NG50 | 89,582,900 | 70,291,102 | 25,168,213 | 21,495,909 | 27,796,056 |
|  | NA50 | 83,951,769 | 67,915,433 | 25,210,265 | 22,128,410 | 28,254,983 |
|  | NGA50 | 83,951,769 | 70,291,086 | 25,167,913 | 21,495,872 | 27,796,056 |
|  | # Misassemblies | 83 | 36 | 224 | 10 | 8 |
|  | Genome fraction (%) | 97.479 | 97.667 | 95.471 | 96.351 | 96.670 |

<sup>\*</sup>MP: multipass mode (**--multipass**), 16 threads, 7.01 min, 10.67 GB peak memory. We did not run Gallifrey-MP on *D. melanogaster* data (—).

We include Hifiasm and HiCanu as state-of-the-art references for assembly accuracy and contiguity. Our primary competitor is rust-mdbg, which operates in a similar speed and contiguity regime to Gallifrey.

For *D. melanogaster*, Gallifrey has lower N50 and NG50 values than the other tools, but produces substantially fewer misassemblies than all of them, including rust-mdbg. The lower contiguity is partly attributable to Gallifrey’s conservative assembly, in which longer contigs spanning difficult regions are broken or excluded rather than assembled with errors. To assess this, we removed QUAST-identified misassembled contigs and re-evaluated the remaining contigs. After filtering, Gallifrey retains a substantially higher genome fraction than rust-mdbg (84.215% vs. 43.353%), with an NG50 of 1.37 Mb compared to only 12 Kbp for rust-mdbg (see Table S2). These results indicate that much of rust-mdbg’s assembly coverage is contained in misassembled contigs, whereas Gallifrey retains most of its genome coverage.

For *H. sapiens*, Gallifrey has substantially fewer misassemblies than all other tools, including rust-mdbg, while maintaining a competitive genome fraction. Gallifrey also supports a multipass mode (--multipass) that assembles reads over successive passes with increasing window lengths to address repeats and coverage gaps. Multipass assembly further reduces misassemblies and substantially improves N50 and NG50 (the Gallifrey-MP column), surpassing rust-mdbg in contiguity while retaining its accuracy advantage. At 16 threads, multipass assembly took 7.01 minutes and used 10.67 GB of peak memory.

### 3.2 Performance analysis on a personal computer

We evaluated Gallifrey on 30*×* PacBio HiFi data from CHM13, a haploid human genome cell line, and compared it with rust-mdbg on an Apple M4 MacBook Pro with 12 cores and 24 GB of memory. To our knowledge, rust-mdbg is the only other HiFi assembler that can run on a PC. Therefore, we limited the comparison between rust-mdbg and Gallifrey.

Compared with the complete rust-mdbg, Gallifrey achieved an 8.1*×* reduction in wall-clock time (58.2 vs. 470 s) and used approximately half the peak memory (5.38 vs. 10.78 GB). Additionally, Gallifrey required no temporary disk space, whereas rust-mdbg used 20.6 GB (Table S3).

## 4 Discussion and Future work

In this work, we introduced Gallifrey, a long-read de Bruijn graph assembler that replaces minimizer sketching with Locally Consistent Seed Clouds (LCSC). By anchoring multi-level parsing on context-free open syncmers and maintaining uncompressed sequence deltas, Gallifrey reconciles the bounded-gap guarantees and near-uniform spacing of LCP with the throughput required for whole-genome datasets. Furthermore, our strand-reconciliation pipeline, which couples two-hit Bloom filter gating, retro-admission, and core-space bipartite graph layouts, effectively overcomes the inherent chirality of deterministic parsing. On standard benchmarks, Gallifrey assembles a human genome in under one minute on commodity hardware while reducing structural misassemblies by up to an order of magnitude compared to rust-mdbg.

Currently, Gallifrey focuses on ultra-fast draft assembly and omits read-level error consensus, base polishing, and haplotype phasing. While our multipass heuristic partially addresses repeats and coverage dropouts, sketch-space resolution still limits the deconvolution of complex nested duplications. Future work will build upon Gallifrey’s underlying primitives: the bipartite LCSC overlap engine naturally extends to rapid sequence-to-graph mapping, while lightweight consensus and phasing could be performed directly in core space. Furthermore, Gallifrey’s ability to efficiently represent dense seed spaces (e.g., syncmers) alongside the bounded perturbation and predictable locality of LCSC (Lemma 1) provides an ideal foundation for resolving heterozygosity in diploid genomes, profiling diversity in metagenomic binning, and building pan-genome variation graphs directly from raw long reads.

## 5 Acknowledgements

## LLM Usage Statement

Claude Opus and Gemini Argon were used to assist in implementing Gallifrey, deriving the lemmas, LATEX typesetting, and correcting the grammar in the manuscript. All generated code and text are inspected, reviewed, and corrected by the authors.

