## Supplemental Text for "Fast draft assemblies using locally consistent parsing"

### Supplementary material

### S1 Bloom filter

A Bloom filter [27] is a space-efficient probabilistic set that inserts and tests an element by setting and checking $h$ hashed bit positions, with no false negatives and a tunable false-positive rate. In Gallifrey, it records the lcmer's observed at least once, so that an lcmer enters the dBG only on its second occurrence (Algorithm S2). Lcmers arising from sequencing errors typically occur once and are thus excluded without being stored; a false positive may admit such an lcmer, but never removes a genuine one. A standard Bloom filter scales poorly in our application: each operation touches $h$ random positions in a bit array far larger than the CPU caches, and concurrent insertions from many threads must be synchronized. Gallifrey therefore uses a blocked Bloom filter [25], in which one hash value maps each element to a single (64-byte) block, the size of a cache line, and all of its bits are set and tested within that block, at the cost of a slightly higher false-positive rate for a given filter size. Each block is stored as eight 64-bit atomic words: an insertion applies a relaxed atomic fetch-or to only the words that lack one of its bits, and a query uses relaxed atomic loads, so threads insert concurrently without locks.

The filter is configured by a target false-positive rate $\varepsilon$ and a projected number $n$ of distinct lcmers, with defaults $\varepsilon = 10^{-3}$ and $n = 10^8$. Each element uses $\mathrm{round}(\log_2(1/\varepsilon))$ hash values: one selects its block, and the remaining $h$ select bit positions drawn independently and uniformly from the 512 bits of that block ($h = 9$ for $\varepsilon = 10^{-3}$). Because blocks receive unequal numbers of elements, the standard Bloom filter formula underestimates the false-positive rate of a blocked filter, so we size the filter with an exact model instead. With $b$ blocks, the number of elements in a block is Poisson-distributed with mean $\mu = n/b$; a block holding $j$ elements has the number of set bits $s$ obtained from $jh$ uniform draws over its 512 bits, and a new element mapped to it is a false positive with probability $(s/512)^h$. The filter uses the smallest $b$ for which this modelled rate is at most $\varepsilon$.

**Algorithm S1** lcmer keys of a read. Arithmetic is modulo $2^{32}$, and each key is $hi \cdot 2^{32} + lo$.

**Require:** cores $\rho[0 \dots n-1]$ with $n \geq W + 2$; polynomial base $B$

$hi \leftarrow 0$
**for** $i \leftarrow 1$ **to** $W - 1$ **do** ▷ $\rho[0]$ is skipped
    $hi \leftarrow hi \cdot B + \rho[i]$
**end for**
$lo \leftarrow \rho[W]$
**emit** $(hi, lo)$ ▷ window $\rho[1 \dots W]$
**for** $i \leftarrow W + 1$ **to** $n - 2$ **do** ▷ $\rho[n-1]$ is skipped
    $hi \leftarrow (hi - \rho[i - W] \cdot B^{W-2}) \cdot B + lo$ ▷ drop oldest core, shift in old last core
    $lo \leftarrow \rho[i]$
    **emit** $(hi, lo)$ ▷ window $\rho[i - W + 1 \dots i]$
**end for**

**Algorithm S2** Graph construction. lcmer keys are computed by Algorithm S1.

**Require:** reads $R$; window length $W$; Bloom filter $BF$; lcDBG $G$ (key → node)
**Require:** $\bar{k}$: key $k$ with a strand tag, recording a reverse-complement sighting

```
for all r ∈ R in parallel do
    K[1...m] ← lcmer keys of r (Algorithm S1); skip r if it has fewer than W + 2 cores
    for i ← 1 to m do
        adm[i] ← BF.TEST_AND_INSERT(K[i]) or BF.TEST(K̄[i])                ▷ seen on either strand
    end for
    for i ← 1 to m do                                           ▷ adm[0] = adm[m + 1] = false
        if adm[i] and (adm[i − 1] or adm[i + 1]) then
            if K[i] ∉ G then G[K[i]] ← NODE(fragment_i(r))
            if adm[i − 1] then ADD_EDGE(K[i − 1] → K[i])
            if adm[i + 1] then G[K[i]].count += 1
        end if
    end for
    L[1...m′] ← lcmer keys of revcomp(r)                                          ▷ priming
    for j ← 1 to m′ do
        fresh[j] ← BF.TEST(L[j]) and not BF.TEST_AND_INSERT(L̄[j])
        if fresh[j] then                                ▷ all earlier sightings own-strand
            if L[j] ∉ G then G[L[j]] ← NODE(fragment of L[j] in revcomp(r))
            G[L[j]].owes ← true
        end if
        if j > 1 and L[j − 1], L[j] ∈ G and (G[L[j − 1]].owes or G[L[j]].owes) then
            if fresh[j − 1] or fresh[j] then                              ▷ retro admission
                ADD_EDGE(L[j − 1] → L[j]); G[L[j − 1]].count += 1
            else if L[j − 1] → L[j] ∉ G then
                OFFER(L[j − 1] → L[j])
            end if
        end if
    end for
end for
for all offered u → v do                                  ▷ decided before any is added
    if u ≠ v, u has no successor, and v has no predecessor then
        if v → u is not offered, u is offered no other successor, and v no other predecessor then
            ADD_EDGE(u → v); G[u].count += 1
        end if
    end if
end for
remove nodes with count = 0 and no incoming edge
```

**Algorithm S3** Parallel simple-path merging.

```
Require: lcDBG $G$; each node $v$ holds its predecessor list $pred(v)$ and its number of successors $outdeg(v)$
$Z \leftarrow \emptyset$                                                    ▷ last nodes of simple paths
for all $v \in G$ in parallel do                                            ▷ each thread scans its own shards
    if $|pred(v)| = 1$ and $outdeg(v) \neq 1$ then
        add $v$ to $Z$
    else if $|pred(v)| > 1$ then
        for all $p \in pred(v)$ with $p \neq v$, $|pred(p)| = 1$ and $outdeg(p) = 1$ do
            add $p$ to $Z$
        end for
    end if
end for
for all $z \in Z$ in parallel do                                            ▷ paths are node-disjoint; no locks
    $u \leftarrow z$; $P \leftarrow [\,]$
    while $pred(u) = \{p\}$ with $p \neq u$ and $outdeg(p) = 1$ do          ▷ trace backward
        append $u$ to $P$; $u \leftarrow p$
    end while
    if $P \neq [\,]$ then
        append the fragments of $P$ to $u$ in path order                    ▷ $P$ reversed
        $count(u) \leftarrow count(u) + \sum_{x \in P} count(x)$
        $outdeg(u) \leftarrow outdeg(z)$
        for all $x \in P$ do redirect $x \to u$
    end if
end for
rewrite every predecessor entry through the redirects; drop the self-edges this creates
```

**Algorithm S4** Tip removal and bubble popping.

```
Require: lcDBG G; tip depth τ; maximum tip length T; locus depth limit d_max; length ratio bound β
procedure CUT_TIPS(G, τ)
    for all v ∈ G do
        if depth(v) < τ then
            if indeg(v) = 0 and outdeg(v) = 0 then remove v                          ▷ isolated
            else if indeg(v) = 1 and outdeg(v) = 0 then remove v                     ▷ dead end
        else
            for all p ∈ pred(v) with indeg(p) = 0 and outdeg(p) = 1 do     ▷ sources entering v
                if depth(p) < τ then remove p
                else if depth(p) < depth(v) and len(p) < T and len(v) > 2T then remove p
            end for
        end if
    end for
end procedure

procedure POP_BUBBLES(G, d_max, β)
    for all s with outdeg(s) ≥ 2, smallest bubble first do
        (t, I) ← SINK(s) in the current graph; if it fails then continue
        p ← heaviest s → t path through I by summed counts
        R ← I \ p                                                          ▷ nodes to remove
        if R = ∅ or len(R) > β · max(1, len(p)) then continue
        if depth(x) ≥ d_max for any x ∈ {s, t} ∪ I then continue   ▷ a repeat: the arms are different copies
        remove R and every edge among {s, t} ∪ I that is not on s → p → t
    end for
end procedure

procedure SINK(s)                                                      ▷ as in miniasm [30]
    ready ← [s]; visited ← ∅; reached ← ∅; I ← [ ]
    while ready ≠ [ ] do
        v ← pop(ready); add v to visited; remove v from reached
        if v has a self-loop then fail
        if v ≠ s then append v to I
        if |I| > 256 or len(I) > 1024 then fail
        if succ(v) = ∅ then fail                                ▷ dead end inside the bubble
        for all w ∈ succ(v) do
            if w = s then fail                                          ▷ a cycle through s
            add w to reached
            if pred(w) ⊆ visited then push w to ready
        end for
        if |ready| = 1 and |reached| = 1 then
            t ← the node in ready
            if I = [ ] or t has a self-loop or an edge to s then fail
            return (t, I)
        end if
    end while
    fail
end procedure
```

**Algorithm S5** Merging the unitigs of a layout into segments.

**Require:** unitigs $u_1, \ldots, u_n$ of one layout, sorted by position, each with its assigned orientation and position
**Require:** shift tolerance $\delta$ (in cores)

**if** the layout has at most one unitig per orientation **then**
  **return** the longer unitig
**end if**
$D \leftarrow$ chain of the cores of $u_1$, each node with weight 1
**for** $j \leftarrow 2$ to $n$ **do**
  $\rho[1 \ldots M] \leftarrow$ LCSC cores of $u_j$ in its assigned orientation
  $S \leftarrow \{(v, i) : v \in D,\ \text{label}(v) = \text{label}(\rho[i]),\ \text{hpc}(v) = \text{hpc}(\rho[i])\}$
  $C \leftarrow$ longest subsequence of $S$ increasing in both the topological order of $v$ and $i$
  split $C$ where the layout shift $\text{pos}(\rho[i]) - \text{pos}(v)$ changes by more than $\delta$; drop pieces with a single match
  **for** $i \leftarrow 1$ to $M$ **do**
    **if** $(v, i) \in C$ for some $v$ **then**
      $x_i \leftarrow v$; $\text{weight}(v) \leftarrow \text{weight}(v) + 1$
    **else**
      $x_i \leftarrow$ new node for $\rho[i]$ with weight 1
    **end if**
    **if** $i > 1$ **then** add edge $x_{i-1} \to x_i$
    **end if**
  **end for**
**end for**
**for** each weakly connected component $D_k$ of $D$ **do**
  emit the heaviest path of $D_k$ as a segment, taking the bases of each core from one unitig that shares it
**end for**

#### S1.1 General tool parameters for the server experiments

For `rust-mdbg`, we used the same parameters as reported in the original publication [14] for both the *D. melanogaster* and HG002 dataset. Similarly, HiCanu and Hifiasm were run with the default parameters recommended in their respective publications [4,5].

**Table S1.** Parameters used for each tool and dataset on the server.

| Tool | Dataset | Parameters |
|---|---|---|
| `rust-mdbg` | *D. mel.* | `-k 35 -l 12 --density 0.002 --bf` |
| | CHM13 | `-k 21 -l 14 --density 0.003 --bf` |
| GALLIFREY | *D. mel.* | `--syncmer 12/3/4 --window 8 -l 6` |
| | CHM13 | `--syncmer 12/4/5 --window 8 -l 5` |
| GALLIFREY-MP | CHM13 | `--syncmer 12/4/5 --multipass 4:8,5:10` |
| QUAST | — | `--fast --large --split-scaffolds` |

### S2 Theoretical properties of LCSC

*Setting.* Let $\tilde{S} = \text{HPC}(S)$ be a homopolymer-compressed sequence of length $n$. At level 1, Gallifrey selects open syncmers [23] with parameters $(k, s, t)$, $0 \leq t \leq k - s$. A $k$-mer is selected if and only if the leftmost occurrence of its smallest $s$-mer lies at offset $t$. Let $\mathcal{P}^{(1)} = (\rho_1^{(1)}, \ldots, \rho_{m_1}^{(1)})$ be the resulting vector of level-1 seed-cores, ordered by position.

For each level $\ell \in \{2, \ldots, L\}$, LCSC applies one round of Deterministic Coin Tossing (DCT) to $\mathcal{P}^{(\ell-1)}$. It then applies the four LCP rules (LMIN, LMAX, RINT, SSEQ) [21] to the reduced sequence, which yields $\mathcal{P}^{(\ell)}$. A level-$\ell$ core $x$ is the concatenation of consecutive level-$(\ell - 1)$ cores $y, z, w, \ldots$, and its label is $x'' = \psi(y'', z'', w'', \ldots)$, computed with MurmurHash [21]. For each level $\ell$, let $m_\ell = |\mathcal{P}^{(\ell)}|$ denote the number of level-$\ell$ seed-cores; in particular, $m_1$ is the number of selected seeds. We write $c_\ell = m_{\ell-1}/m_\ell$ for the reduction factor at level $\ell$.

We use two properties of LCP proven in [21]: *Contiguity*, meaning there are no gaps between consecutive cores; and *Adjacency*, meaning at most two of any three consecutive symbols start a core. Both carry over to LCSC unchanged, because LCSC applies the same rules, only to a different input alphabet.

*Remark S1 (Alphabet Independence of the Adjacency Property).* While the Adjacency Property ($m_\ell \leq \frac{2}{3} m_{\ell-1} + O(1)$) was established over the four-letter DNA alphabet [21], the property holds unconditionally for all hierarchy levels $\ell \geq 2$ regardless of alphabet expansion [20]. The four LCP partitioning rules rely strictly on pairwise equality ($=$) and the strict total order ($<$) of adjacent symbols [20]. Because Deterministic Coin Tossing maps level-$(\ell - 1)$ core identifiers to fixed-width bitstrings endowed with standard lexicographical comparison, the reduced symbol alphabet $\Sigma_\ell$ forms a valid totally ordered domain [20]. Consequently, no three consecutive symbols in the reduced sequence can yield three overlapping core starts, preserving the worst-case bound $c_\ell \geq 3/2$ across all levels [20].

#### Core counts and parsing time

**Proposition S1 (Geometric core reduction).** *Assume the $s$-mers within each $k$-mer are distinct, as in the standard random-sequence model. Then the following hold.*

1. $\mathbb{E}[m_1] = n/(k - s + 1)$.
2. $m_L = m_1 / \prod_{\ell=2}^{L} c_\ell$, *with* $c_\ell \geq 3/2$ *up to an additive* $O(1)$ *boundary term. In the worst case, therefore,* $m_L \leq (2/3)^{L-1} m_1 + O(1)$.
3. *The mean distance between the starts of consecutive level-$L$ cores is* $n/m_L$, *which equals* $(k - s + 1) \prod_{\ell=2}^{L} c_\ell$ *in expectation.*
4. *Seed selection takes* $O(n)$ *time, and building all levels* $2, \ldots, L$ *takes* $O(m_1)$ *time.*

*Proof.* (1) Under the distinctness assumption, the leftmost minimum of the $k - s + 1$ $s$-mers in a $k$-mer is equally likely to lie at any offset. A $k$-mer is therefore selected with probability $1/(k-s+1)$, and the claim follows by linearity of expectation.

(2) The identity holds by the definition of $c_\ell$. For the bound, apply Adjacency Property to the reduced sequence at level $\ell - 1$, which has length $m_{\ell-1}$. At most $2\lceil m_{\ell-1}/3 \rceil$ of its positions start a core, so $m_\ell \leq \frac{2}{3} m_{\ell-1} + O(1)$.

(3) This follows from (1) and (2).

(4) Open syncmers can be selected in $O(n)$ time with a sliding-window minimum over $s$-mers. Level $\ell$ is computed in $O(m_{\ell-1})$ time, because DCT and each LCP rule inspect a constant number

of neighbors. Summing over levels gives

$$\sum_{\ell=1}^{L-1} m_\ell \;\le\; m_1 \sum_{j\ge 0} (2/3)^j + O(L) \;=\; 3m_1 + O(L).$$

By the Contiguity Property, every level-1 seed is covered by some level-$L$ core. The exceptions are seeds near the ends of the vector and the rare short stranded regions that a single DCT round can leave uncovered [21].

**Empirical grounding** On CHM13v2.0, nucleotide-level LCP has a reduction factor of $c_\ell \approx 2.34$ at every level $\ell \ge 2$ [21]. This is well above the worst-case value of 3/2. Applied to raw nucleotides, LCP produces $1.39 \times 10^9$ level-1 cores from a single genome copy [21], so a 30× HiFi dataset would yield on the order of $4 \times 10^{10}$ level-1 cores.

LCSC instead starts from open syncmers ($k = 12$, $s = 4$, $t = 5$, density 1/9) and parses up to $L = 5$. Assuming the same reduction factor, this gives an expected sparsification of $(k-s+1)\, c^{L-1} \approx 9 \times 2.34^4 \approx 270$ relative to the homopolymer-compressed bases. For comparison, nucleotide-level LCP has a coefficient of variation of 0.28–0.30 for inter-core distance at levels 5–8 [21].

**Locality of perturbations** For two core vectors that agree outside one contiguous block, we write

$$C = A\,X\,B, \qquad C' = A\,X'\,B,$$

where $A$ and $B$ are shared. We call $\Delta = \max(|X|, |X'|)$ the size of the differing block. Throughout, we assume the edit lies far enough from both ends of the sequence that boundary effects do not reach it.

**Lemma 1 (restated).** *Let $\tilde{S}$ and $\tilde{S}'$ differ by a single-character edit, and let $d = \min(t, k-s-t)+1$. Then the following hold. (1) $\Delta_1 \le \lceil k/d \rceil$, and each of $X$ and $X'$ has expected size at most $k/(k-s+1)$. (2) $\Delta_\ell \le \lceil (\Delta_{\ell-1} + r)/c \rceil$ with $r = 12$ and $c = 3/2$. Moreover, $\Delta_\ell = 0$ whenever $\Delta_{\ell-1} = 0$. (3) The edit changes at most $\Delta_L + W - 1$ consecutive lcmers on each side.*

**Lemma S1 (Syncmer spacing).** *Any two selected open syncmers start at least $d = \min(t, k-s-t)+1$ positions apart.*

*Proof.* Let $\sigma_j$ denote the $s$-mer starting at position $j$. Suppose the $k$-mers starting at $i < j$ are both selected, and that $j - i \le \min(t, k-s-t)$.

Because $j - i \le k - s - t$, position $j + t$ lies in the $s$-mer window $[i,\, i+k-s]$ of the $k$-mer at $i$. That $k$-mer is selected, so its leftmost minimum is at $i + t$, which gives $\sigma_{j+t} \ge \sigma_{i+t}$.

Because $j - i \le t$, position $i + t$ lies in the window $[j,\, j+k-s]$ of the $k$-mer at $j$, and $i + t$ precedes $j + t$. Since the leftmost minimum of that $k$-mer is at $j + t$, we get $\sigma_{i+t} > \sigma_{j+t}$.

The two inequalities contradict each other.

*Proof (Proof of Lemma 1(1)).* A $k$-mer's selection status and its label depend only on its own bases. For a substitution at position $p$, the $k$-mers covering $p$ start in the window $[p-k+1,\, p]$. For an insertion or a deletion, the $k$-mers touching the edit in each string likewise start within at most $k$ consecutive positions.

Seeds that start outside this window are identical in both strings and appear in the same order. They form $A$ and $B$, and the selected seeds inside the window form $X$ and $X'$. By Lemma S1, at most $\lceil k/d \rceil$ selected seeds start within $k$ consecutive positions, which bounds both $|X|$ and $|X'|$.

For the expectation, each of the at most $k$ positions in the window is selected with probability $1/(k-s+1)$ (Proposition S1).

For the default parameters, $k=12$ and $s=4$, so $k-s=8$. With $t=5$ (or $t=4$ under 0-based indexing), $d \geq 4$, and therefore $\Delta_1 \leq 3$.

Note that $X = X' = \emptyset$ is possible. An edit in a base that no selected seed covers changes no core at any level. In that case the edit affects only the sequence fragment stored in a node (Section 2.5).

**Lemma S2 (One-level locality).** *Suppose* $C^{(\ell-1)} = A\,X\,B$ *and* $C'^{(\ell-1)} = A\,X'\,B$, *with* $|X|, |X'| \leq \Delta$. *Then* $C^{(\ell)}$ *and* $C'^{(\ell)}$ *also decompose as shared prefix, differing block, shared suffix, and each differing block has at most* $\frac{2}{3}\Delta + 8$ *cores. If* $\Delta = 0$, *the two vectors are identical.*

*Proof.* The proof has four steps.

*(i) DCT.* The reduced code of core $j$ depends only on the bitstreams $b_{j-1}$ and $b_j$. If $b_{j-1} = b_j$, we assume DCT emits a fixed sentinel code, which is still a function of the pair. The reduced sequences therefore agree everywhere except on the symbols of $X$ (or $X'$) and on the first symbol of $B$. Call this differing block $Y$; it satisfies $|Y| \leq \Delta + 1$.

*(ii) Boundaries are decided locally.* Whether a symbol starts or ends a core depends only on symbols within distance 3 of it:

- LMIN inspects $xyz$.
- LMAX inspects $sxyzt$.
- The start and the end of a RINT core $xy^iz$ are fixed by the pattern $x \neq y = y$ and by $y = y \neq z$, respectively.
- An SSEQ core starts at the last symbol of a flanking core, which takes LMAX's reach of 3, and it ends where the next such core starts.

*(iii) Labels.* The label of a level-$\ell$ core is $\psi$ applied to the labels of its constituent cores. It is therefore the same in both strings whenever all its constituents lie in $A$, or all lie in $B$.

*(iv) Counting.* By (ii) and (iii), a level-$\ell$ core can differ between the two strings only if its extent meets $I = Y$ extended by 3 symbols on each side, so $|I| \leq \Delta + 7$. By Contiguity, the cores meeting $I$ are consecutive in the vector, which gives the claimed decomposition. By Adjacency, at most $2\lceil |I|/3 \rceil \leq \frac{2}{3}|I| + \frac{4}{3}$ cores start inside $I$. Consecutive cores overlap by at most two symbols, so at most two cores that start before $I$ extend into it. In total, each block has at most

$$\tfrac{2}{3}(\Delta + 7) + \tfrac{4}{3} + 2 \;=\; \tfrac{2}{3}\Delta + 8$$

cores.

If $\Delta = 0$, then $Y$ is empty, and every core coincides in the two strings by (ii) and (iii).

*Proof (Proof of Lemma 1(2)).* Lemma S2 gives

$$\Delta_\ell \;\leq\; \tfrac{2}{3}\big(\Delta_{\ell-1} + 12\big) \;=\; \frac{\Delta_{\ell-1} + r}{\rho}$$

with $r = 12$ and $c = 3/2$. The map $x \mapsto \frac{2}{3}x + 8$ is a contraction with fixed point 24. Since $\Delta_1 \leq 3 < 24$, we have $\Delta_\ell \leq 24$ for every $\ell$, independently of $n$ and $L$. This is consistent with the main-text bound $(r+c)/(c-1) = 24$, which also absorbs the ceiling.

These constants are worst-case bounds. With the empirical reduction factor $c \approx 2.34$, the fixed point drops to about 9, and the typical block is much smaller (Fig. 1c).

*Proof (Proof of Lemma 1(3)).* An lcmer is a window of $W$ consecutive level-$L$ cores, and its key is a function of those cores (Algorithm 1). Write $C^{(L)} = A\,X\,B$ and $C'^{(L)} = A\,X'\,B$. Windows that lie entirely within $A$, or entirely within $B$, have the same key in both strings. A window

differs only if it meets $X$ (respectively $X'$), and there are $|X|+W-1$ (respectively $|X'|+W-1$) such windows.

Assume there are no key collisions and that these lcmers do not occur elsewhere in the genome. The two reads then share lcDBG nodes up to the last window inside $A$ and again from the first window inside $B$. Between these shared nodes they follow disjoint arms of $|X|+W-1$ and $|X'|+W-1$ nodes, which form a bubble. For a substitution with $|X| = |X'| = 1$, each arm has exactly $W$ nodes. An indel can produce arms of unequal length.

#### S2.1 Aditional QUAST experiment for *D. melanogaster*

**Table S2.** QUAST assembly quality metrics for *D. melanogaster* after filtering misassembled contigs.

| **Metric** | **rust-mdbg** | **Gallifrey** |
|---|---:|---:|
| N50 | 8,242,452 | 1,473,341 |
| NG50 | 12,072 | 1,368,453 |
| NA50 | 8,232,638 | 1,473,341 |
| NGA50 | 9,058 | 1,368,453 |
| # Misassemblies | 0 | 0 |
| Genome fraction (%) | 43.353 | 84.215 |

*Both tools run at 16 threads. Misassembled contigs were identified using QUAST and removed prior to re-evaluation.*

**Table S3.** Run time and resources on a laptop.

| **Tool** | **Wall time (s)** | **CPU time (s)** | **Peak memory (GB)** | **Temporary disk (GB)** |
|---|---:|---:|---:|---:|
| Gallifrey | 58.2 ± 0.8 | 492 ± 2 | 5.38 ± 0.01 | 0 |
| `rust-mdbg` (Total) | 470 ± 3 | 1530 ± 12 | 10.78 ± 0.06 | 20.6 |
| `rust-mdbg` | 267 ± 3 | 1332 ± 12 | 10.78 ± 0.06 | 11.2 |
| `magic_simplify` | 203 ± 1 | 195 ± 1 | – | 9.4 |

32× coverage CHM13 HiFi reads assembled on an Apple M4 Pro with 12 threads. Values are the mean ± s.d. of three runs.

### S3 Graph simplification details

During construction, Gallifrey stores only the incoming edges of each node, as a list of its predecessors. The initial merge of simple nodes needs only this list and the number of outgoing edges of each node, which avoids storing edge information twice while the graph is at its largest. Before this merge, we count the outgoing edges of every node by traversing the graph once. After the initial merge, the graph is much smaller, and we compute the outgoing edges of the remaining nodes from their incoming edges for the steps that follow edges forward.

We refer to nodes that have a single incoming and a single outgoing edge as simple nodes. We simplify the graph by merging consecutive simple nodes, removing tips and popping bubbles. As the latter two operations may create new simple nodes, we repeat this procedure until convergence.

**Algorithm S6** Graph cleaning.

**Require:** lcDBG $G$; maximum rounds $N$; length ratio bound $\beta$

MERGE_SIMPLE_PATHS($G$) ▷ Algorithm S3
remove every $v$ with $indeg(v) = outdeg(v) = 1$ and $depth(v) < 2$
$d_1 \leftarrow$ lowest substantial depth peak of long nodes ▷ one-copy depth
$d_{\max} \leftarrow \tilde{d}\,(1 + 3\,Cov_v)$ of fork depths near $d_1$ ▷ locus depth limit
$\tau \leftarrow \max(1, \mathrm{round}(d_1/2))$ ▷ tip depth
**while** $G$ has not converged **and** $round < N$ **do**
    CUT_TIPS($G, \tau$)
    POP_BUBBLES($G, d_{\max}, \beta$) ▷ Algorithm S4
    MERGE_SIMPLE_PATHS($G$)
**end while**
**return** $G, d_{\max}$ ▷ $d_{\max}$ is reused by dead-end joining (Section 2.8)

**Lock-Free Parallel Path Contraction** Since most nodes are simple after construction, the initial merge of simple nodes is the most expensive step, and we devised a parallel algorithm for it (Algorithm S3). The dBG is stored in a sharded concurrent hash table, where graph operations acquire a lock only on the shard containing the node being operated on. A merge, however, involves several nodes that are unlikely to lie in the same shard, so naive locking is prone to deadlocks. We avoid locking altogether. Because nodes store their incoming edges, each path of simple nodes is merged backward, starting from its last node. This node can be recognized from itself and its neighbors alone: it has a single incoming edge, and either it has zero or several outgoing edges, or its successor has several incoming edges. Each path has exactly one last node, so every simple node belongs to exactly one path and no two paths share a node. The only exception is a subgraph that is entirely circular, in which every node is simple; it has no last node and is left unmerged. We therefore find all last nodes in parallel, with each thread scanning separate shards of the dBG, and then merge their paths in parallel without acquiring the shard locks. Since the paths do not share nodes, no race conditions arise.

**Depth thresholds** The depth of a node is its count divided by its length in lcmers, that is, the mean read support of its lcmers. After the initial contraction, we remove nodes with a single incoming and a single outgoing edge and a depth below 2. The one-copy depth $d_1$ is the lowest substantial peak of the depth distribution of long nodes (Figure S1.a). Collapsed repeats only add depth, so single-copy sequence forms the lowest peak. The locus depth limit is $d_{\max} = \tilde{d}\,(1+3Cov_v)$, where $\tilde{d}$ is the median depth of nodes with multiple outgoing edges and depth between $d_1/2$ and

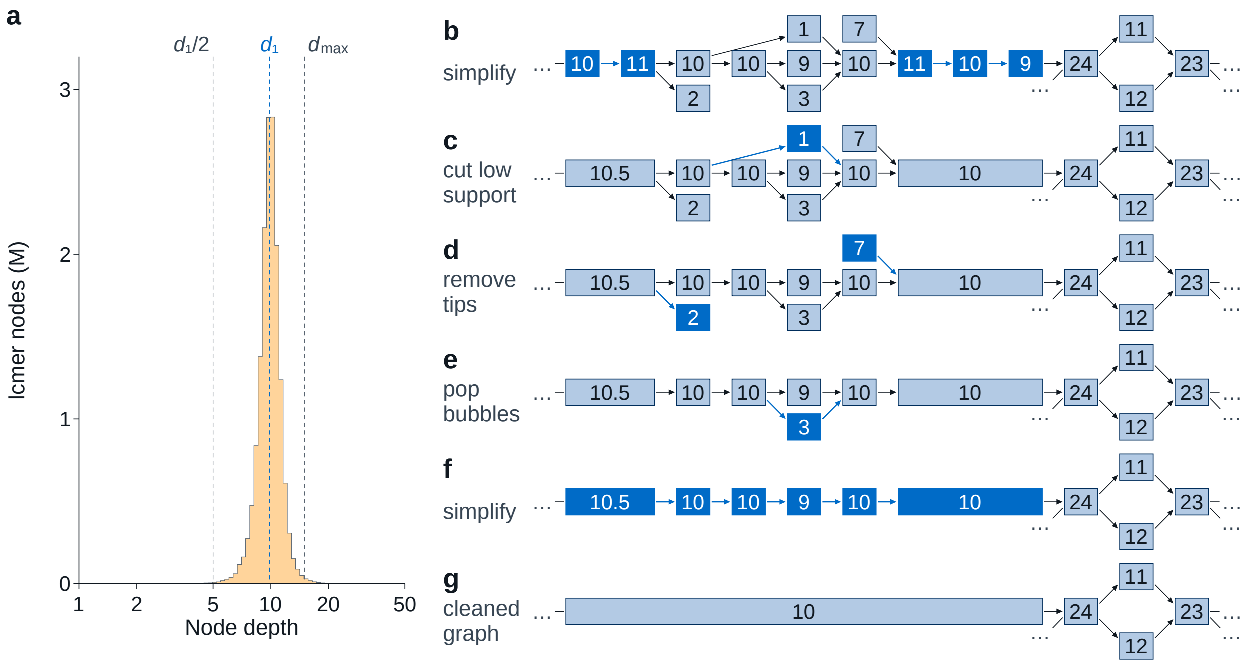


**Fig. S1. Graph cleaning.** (a) Depth distribution of long nodes after the initial simplification, with the one-copy depth $d_1$, the tip depth $d_1/2$ and the locus depth limit $d_{\max}$. (b–g) One round of cleaning on a small lcDBG; numbers on the nodes represent the lcmer depths.

$3d_1/2$, and $Cov_v$ is the robust coefficient of variation of these depths. A node deeper than $d_{\max}$ likely holds more than one copy of its sequence.

**Tip removal** Tips are removed with the tip depth $\tau = \max(1, \text{round}(d_1/2))$ (Algorithm S6, Figure S1.d); the procedure is given in Algorithm S4. A node whose depth is below $\tau$ is removed if it is isolated or a dead end, that is, if it has no outgoing edge and at most one incoming edge. For every other node $v$, we examine the sources entering it, namely predecessors with no incoming edge and a single outgoing edge. Such a source is removed if its depth is below $\tau$, or if it is shallower than $v$ and shorter than the maximum tip length $T$ while $v$ is longer than $2T$.

**Bubble popping** Bubbles are detected from each node $s$ with at least two outgoing edges, with a superbubble-style search as in miniasm [30] (Algorithm S4, Figure S1.e). The search visits a node only after all of its predecessors have been visited, and reports a sink $t$ when a single node remains to be explored, together with the interior nodes $I$ between $s$ and $t$. It fails if it encounters a self-loop, a dead end, a cycle through $s$, or an interior larger than a fixed limit. Bubbles are processed in order from smallest to largest, and each search runs on the current graph, so that a bubble changed by an earlier pop is evaluated again. Within a bubble, we keep the heaviest path $p$ from $s$ to $t$, measured by the summed counts of its nodes, and remove the remaining interior nodes together with the edges that are not on this path. A bubble is popped only if the removed nodes are at most $\beta$ times as long as $p$ and no node of the bubble reaches the locus depth limit $d_{\max}$; deeper bubbles typically arise from repeats, whose branches are different copies rather than errors.
